# Identification of Genomic Regions Carrying a Causal Mutation in Unordered Genomes

**DOI:** 10.1101/026856

**Authors:** Pilar Corredor-Moreno, Ed Chalstrey, Carlos A. Lugo, Dan MacLean

## Abstract

Whole genome sequencing using high-throughput sequencing (HTS) technologies offers powerful opportunities to study genetic variation. Mapping the mutations responsible for different phenotypes is generally an involved and time-consuming process so researchers have developed user-friendly tools for mapping-by-sequencing, yet they are not applicable to organisms with non-sequenced genomes. We introduce SDM (SNP Distribution Method), a reference independent method for rapid discovery of mutagen-induced mutations in typical forward genetic screens. SDM aims to order a disordered collection of HTS reads or contigs such that the fragment carrying the causative mutation can be identified. SDM uses typical distributions of homozygous SNPs that are linked to a phenotype-altering SNP in a non-recombinant region as a model to order the fragments. To implement and test SDM, we created model genomes with an idealised SNP density based on *Arabidopsis thaliana* chromosome 1 and analysed fragments with size distribution similar to reads or contigs assembled from HTS sequencing experiments. SDM groups the contigs by their normalised SNP density and arranges them to maximise the fit to the expected SNP distribution. We tested the procedure in existing datasets by examining SNP distributions in recent out-cross and back-cross experiments in *Arabidopsis thaliana* backgrounds. In all the examples we analysed, homozygous SNPs were normally distributed around the causal mutation. We used the real SNP densities obtained from these experiments to prove the efficiency and accuracy of SDM. The algorithm was able to successfully identify small sized (10-100 kb) genomic regions containing the causative mutation.

## 1. Background

Forward genetic screens are a fundamental strategy to find genes involved in biological pathways in model species. In these screens a population is treated with a mutagen that alters the DNA of individuals in some way, e.g. induction of guanine-to-adenine substitutions using ethylmethane sulfonate (EMS) [1]. Individuals with a phenotype of interest are then isolated from a mutagenised population and a recombinant mapping population is created by back-crossing to the parental line or out-crossing to a polymorphic ecotype [2]. The recombinant population obtained from that cross will segregate for the mutant phenotype and individuals showing the mutant phenotype will carry the causal mutation, even if the genomic location is unknown. The recombination frequency between the causal mutation and nearby genetic markers is low, so the alleles of these linked genetic markers will co-segregate with the phenotype-altering mutation while the remaining unlinked markers segregate randomly in the genome [3].

The analysis of the allele distribution can uncover these low recombinant regions to identify the location of the causal mutation. This genetic analysis is often referred to as bulked segregant analysis (BSA) [4]. Traditional genetic mapping is a work intensive and time consuming process but recent advances in high-throughput sequencing (HTS) have accelerated the identification of mutations underlying mutant phenotypes in forward genetic screens. Several methods such as CandiSNP [2], SHOREmap [5, 6] or NGM [7] based on bulked segregant analysis of F2 progeny have successfully identified mutants in *Arabidopsis thaliana*. However, all these methods depend on an assembled reference genome and cannot be used in species for which a reference genome is not available. Some alternative solutions using reference sequences of related species have been proposed [8, 9], but these require low sequence divergence and high levels of synteny between the mutant reads and the related reference sequence and this has restrained the application of these approaches [3, 10].

Substantial effort is being made to sequence many species but reasonable completion of a sequence remains expensive and time consuming, and fragmented draft genomes present certain limitations in use for mutation mapping in many circumstances. Fast-evolving and repetitive genes such as disease resistance genes [11] might be absent or divergent from draft reference genome assemblies. Also, draft genomes often contain gaps that can frustrate alignments. In the last few years, several reference-free methods for general mutation identification have been proposed [10, 12, 13, 14] to solve the reference sequence restriction, but none have been extended to allow for direct identification of causative mutations [3, 14, 15].

We propose SDM, a fast causative mutant identification method based on contigs or reads that allows the detection of candidate causative SNPs. Instead of relying on a genome comparison, it focuses on the SNP linkage around the causal mutation and analyses the SNP distribution to identify the chromosome area where the putative mutated gene is located. SDM does not rely on previously known genetic markers and can be used on extremely fragmented genome assemblies, even down to the level of long reads.

## 2. Methods

### 2.1. Model genome generation

We used model genomes to develop our mutant identification method. We assigned an idealised SNP distribution to a set of randomly shuffled sequences that imitate contigs assembled from HTS. We created different model genomes based on *Arabidopsis thaliana* chromosome 1 (TAIR10_chr1.fas, 34.9 Mb) from The Arabidopsis Information Resource [16]. Whole chromosome sequences were obtained from ftp://ftp.arabidopsis.org/home/tair/home/tair/Sequences/whole_chromosomes. *Arabidopsis thaliana* makes an ideal model genome due to its small size, a well-described genetic variation and a small content of repeats.

To generate the model genomes, we used the script https://github.com/edwardchalstrey1/fragmented_genome_with_snps/blob/master/create_model_genome.rb. A detailed protocol and the code to recreate the model genomes are available in a GitHub repository at https://github.com/pilarcormo/SNP_distribution_method/tree/master/Small_genomes.

Homozygous SNPs followed a normal distribution (as proven in the section 3.4). The R function rnorm defined by n, mean and sd was used to describe the homozygous SNP distribution. The mean was specified in the middle of the model genome, generating a normal distribution with equally sized tails. The standard deviation (sd) was 2 times the n value. Heterozygous SNPs followed a uniform distribution in the model genomes. The R function runif defined by n, min and max was used to describe the heterozygous SNPs. The min value was fixed to 1 and the max value was the model genome length. For both functions, n varied in each genome to meet the requirement of finding a SNP every 500 bp so that the resolution of high SNP density peak is good even in small genomes.

A minimum contig size is provided as an argument when running the script, and the maximum contig size is obtained by doubling the minimum value. Contig sizes are randomly chosen to be between these 2 values.

We ran `small_model_genome.rb` and generated 1, 3, 5, 7, 11, 13 and 15 Mb genomes with 1 SNP every 500 bp and 2 different contig sizes to create 1300 and 700 contigs in total. We replicated each genome 5 times, making a total of 70 genomes which can be found at https://github.com/pilarcormo/SNP_distribution_method/tree/master/Small_genomes/arabidopsis_datasets/1-15Mb.

Then, we also ran `chr1_model_genome.rb` to use the whole chromosome 1 length to generate larger model genomes. A more realistic SNP density was used for these models (1 SNP every 3000 bp). In this case, 3 contig sizes were employed to create 1000, 2000 and 4000 contigs approximately and we replicated each model 5 times, obtaining 15 more model genomes. Those were deposited at https://github.com/pilarcormo/SNP_distribution_method/tree/master/Small_genomes/arabidopsis_datasets/30Mb under the names chr1_i for 1000 contigs, chr1_A_i for 2000 contigs and chr1_B_i for 4000 contigs genomes. i ranges from 1 to 5 and is used to name the replicates.

We also generated 2 sets of model genomes with a non-centred mean to test SDM filtering step. These genomes were divided into 2000 contigs. They can be found at https://github.com/pilarcormo/SNP_distribution_method/tree/master/Small_genomes/arabidopsis_datasets/30Mb under the names chr1_C_i, which presents a 20% shift to the right, and chr1_E_i, which presents a 20% shift to the left.

The model genomes directories each contain a FASTA file with the correct fragment order, a FASTA file with the randomly shuffled fragments and a VCF file with the homozygous and heterozygous SNP positions. For simplicity, homozygous SNPs are given a fixed Allele Frequency (AF) of 1 and heterozygous SNPs are given an AF of 0.5 in the VCF file.

### 2.2. SDM implementation using model genomes

Fig. 1 shows the SNP Distribution Method (SDM) workflow.

The first step in the pipeline is to calculate the homozygous to heterozygous SNPs ratio per contig. The ratio of homozygous to heterozygous SNPs on a contig c is defined as the sum of all the homozygous SNPs on c plus 1 divided by the sum of all the heterozygous SNPs on c plus 1.

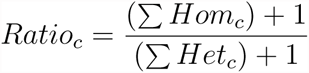

The effect of contig length on SNP density is reduced by normalising the SNP density by length. The absolute number of homozygous SNPs on each contig is divided by the number of nucleotides (contig length) to obtain the contig score:

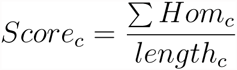

SDM sorts the contigs based on their score so that they follow an ideal normal distribution. It starts by taking the 2 lowest values and locating them at each tail of the distribution. Following this fashion, we obtained the right and left sides that together build up the whole distribution.

The first SDM version was run on the model genomes created as explained in section 2.1. SDM uses as input the VCF file with the homozygous and heterozygous SNP positions, the text files containing the lists of homozygous and heterozygous SNPs and the FASTA file with the shuffled contigs. The FASTA file with the correct contig order is used to calculate the ratios in the correctly ordered fragments so that they can be compared to the ratios obtained after SDM sorts the contigs. The script to run SDM on the model genomes is available at https://github.com/pilarcormo/SNP_distribution_method/blob/master/Small_genomes/SDM.sh.

SDM generates a new FASTA file with the suggested contig order and plots comparing the SNP densities and ratios after SDM to the original values.

For all the 70 genomes ranging from 1 to 15 Mb, no filtering step based on the ratio was used (threshold = 0). The highest kernel density value for the SNP distribution after sorting the contigs with SDM was taken as candidate SNP position. This value was compared to the initial mean of the homozygous SNP distribution to measure the peak shift:

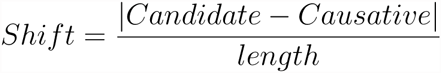

where ‘Candidate’ is the highest kernel density value after SDM and ‘Causative’ is the mean of the normal distribution of homozygous SNPs in the model genome. A CSV file containing all the deviations in the model genomes can be found at https://github.com/pilarcormo/SNP_distribution_method/blob/master/Small_genomes/arabidopsis_datasets/1-15Mb.csv. The same approach was used for the whole-sized genomes (

~~~
chr1_i, chr1_A_i and chr1_B_i
~~~

).

The Ruby code used to run SDM on model genomes is available at the Github repository https://github.com/pilarcormo/SNP_distribution_method/blob/master/Small_genomes/SNP_distribution_method.rb.

### 2.3. SDM deviation in jitter plots

The deviation percentages calculated independently for each genome as described in section 2.2 are available at https://github.com/pilarcormo/SNP_distribution_method/blob/master/Small_genomes/1-15Mb.csv and https://github.com/pilarcormo/SNP_distribution_method/blob/master/Small_genomes/30Mb.csv.

The R code to plot the deviation jitter plots for each genome length and contig size was deposited at https://github.com/pilarcormo/SNP_distribution_method/blob/master/R_cripts/jitter_plots.R.

### 2.4. Pre-filtering step based on the homozygous to heterozygous SNP ratio

The hom/het ratio was used as a cut-off value to discard contigs located furthest away from the causal mutation. If this filtering step is required, the threshold stringency should be provided as an integer. Each integer represents the percentage of the maximum ratio below which a contig will be discarded. For instance, if 1 is specified, SDM will discard those contigs with a ratio falling below 1% of the maximum ratio while a more stringent value of 20 will discard those contigs with a ratio falling below 20% of the maximum ratio.

We used model genomes defined in section 2.1 to test the effectiveness of the filtering step. In particular, we used the model genomes with the normal distribution peak shifted to the right (

~~~
chr1_C_i
~~~

) and the model genomes with the normal distribution peak shifted to the left (

~~~
chr1_E_i
~~~

). Protocol and results were deposited at https://github.com/pilarcormo/SNP_distribution_method/tree/master/Small_genomes/arabidopsis_datasets/Analyse_effect_ratio. The directories 

~~~
chr1_right
~~~

 and 

~~~
chr1_left
~~~

 contain examples of the SDM output after filtering under the names 

~~~
Ratio_0_1
~~~

 (no filtering), 

~~~
Ratio_1_1
~~~

 (1% threshold), 

~~~
Ratio_5_1
~~~

 (5% threshold), 

~~~
Ratio_10_1
~~~

 (10% threshold), 

~~~
Ratio_20_1
~~~

 (20% threshold).

### 2.5. Forward genetic screens used to analyse SNP distribution

We used five different sets of Illumina sequence reads from 4 recent forward genetic screens in *Arabidopsis thaliana* backgrounds [17, 18, 19, 20] (Table 1).

**Table 1.** Forward genetic screens in *Arabidopsis thaliana*, ecotypes involved in the crossing and technologies used to identify the causative mutation, the chromosome (Chr) where the mutation was found and the mutated gene location (Gene) are also specified.

| Sample | Mutant |  | Wild-type | Method | Chr | Gene |
| --- | --- | --- | --- | --- | --- | --- |
| OCF2 | Col-0 | x | Ler-0 | SHOREmap | 2 | SOC1<br>(~18.8 Mb) |
| BCF2 | Col-0 | x | Col-0 | NGM, SHOREmap,<br>GATK and SAMtools | 3 | HASTY<br>(~14.05 Mb) |
| mob1/<br>mob2 | Col-0 | x | Col-0 | CandiSNP | 5 | CPK28<br>(~26.45 Mb) |
| sup#1 | WS | x | Col-0 | Ratio of homozygous<br>to heterozygous SNPs | 4 | SGT1b<br>(~6.85 Mb) |

Galvão et al (**OCF2**) sequenced a mutant pool of 119 F2 mutants generated by out-crossing a Col-0 background mutant to a Ler-0 mapping line. They also sequenced the parental lines and performed conventional SHOREmap [5] to identify a causative mutation [17]. The reads are available to download at http://bioinfo.mpipz.mpg.de/shoremap/examples.html. Allen et al (**BCF2**) back-crossed a Col-0 mutant to the non-mutagenised Col-0 parental line [19]. A pool of 110 mutant individuals showing the mutant phenotype and the parental line were sequenced. They used different SNP identification methods (NGM, SHOREmap, GATK and samtools) [5, 7, 21, 22]. The reads are available to download at http://bioinfo.mpipz.mpg.de/shoremap/examples.html. Monaghan et al obtained two different and independent Col-0 mutants (**bak1-5 mob1** and **bak1-5 mob2**) [20]. The mutants were back-crossed to a parental Col-0 line and sequenced. They used CandiSNP [2] to identify the causal mutation. The last dataset we used (**sup#1**) was obtained by outcrossing an *Arabidopsis Wassilewskija* (WS) mutant to wild-type Col-0 plants followed by sequencing of 88 F2 individuals and WS and Col as parental lines. Uchida et al described a pipeline to identify the causal mutation based on plotting ratios of homozygous SNPs to heterozygous SNPs [18]. Reads are available at http://www.ncbi.nlm.nih.gov/sra/?term=DRA000344.

### 2.6. Read mapping and SNP calling

Mutant and parental reads were subjected to the same variant calling approach. The Rakefile and scripts used to perform the alignment and SNP calling can be found in the Supplementary File 1.

The quality of the deep sequencing was evaluated using FastQC 0.11.2 (http://www.bioinformatics.babraham.ac.uk/projects/fastqc/). Reads were trimmed and quality filtered by Trimmomatic v0.33 [23]. We performed a sliding window trimming, cutting once the average Phred quality fell below 20 in the window size.

The paired-end reads were aligned to the reference sequence of *Arabidopsis thaliana* TAIR10 available from The Arabidopsis Information Resource. The sequence (TAIR10_chr_all.fas) is available at ftp://ftp.arabidopsis.org/home/tair/Genes/TAIR10_genome_release/TAIR10_chromosome_files/ [16]. BWA-MEM long-read alignment using BWA v0.7.5a [24] with default settings was used. The resultant SAM files were converted to BAM files and then sorted using the SAMtools package v1.0 (<u>http://samtools.sourceforge.net/</u>) [22]. Then, we used SAMtools mpileup command to convert the BAM files into pileup files. To call SNPs we used the mpileup2snp command from VarScan v2.3.7 http://varscan.sourceforge.net [25, 26] to get VCF 4.1 output. A default 0.8 allele frequency was used for homozygous SNPs.

VCF files for mutants and mapping lines can be found at the repository https://github.com/pilarcormo/SNP_distribution_method/blob/master/Reads in the individual folder for each screen (OCF2, BCF2, Aw_sup1-2, m_mutants). The **Additional Figure 1** summarises the pipeline used for read mapping and SNP calling.

**Figure 1.**
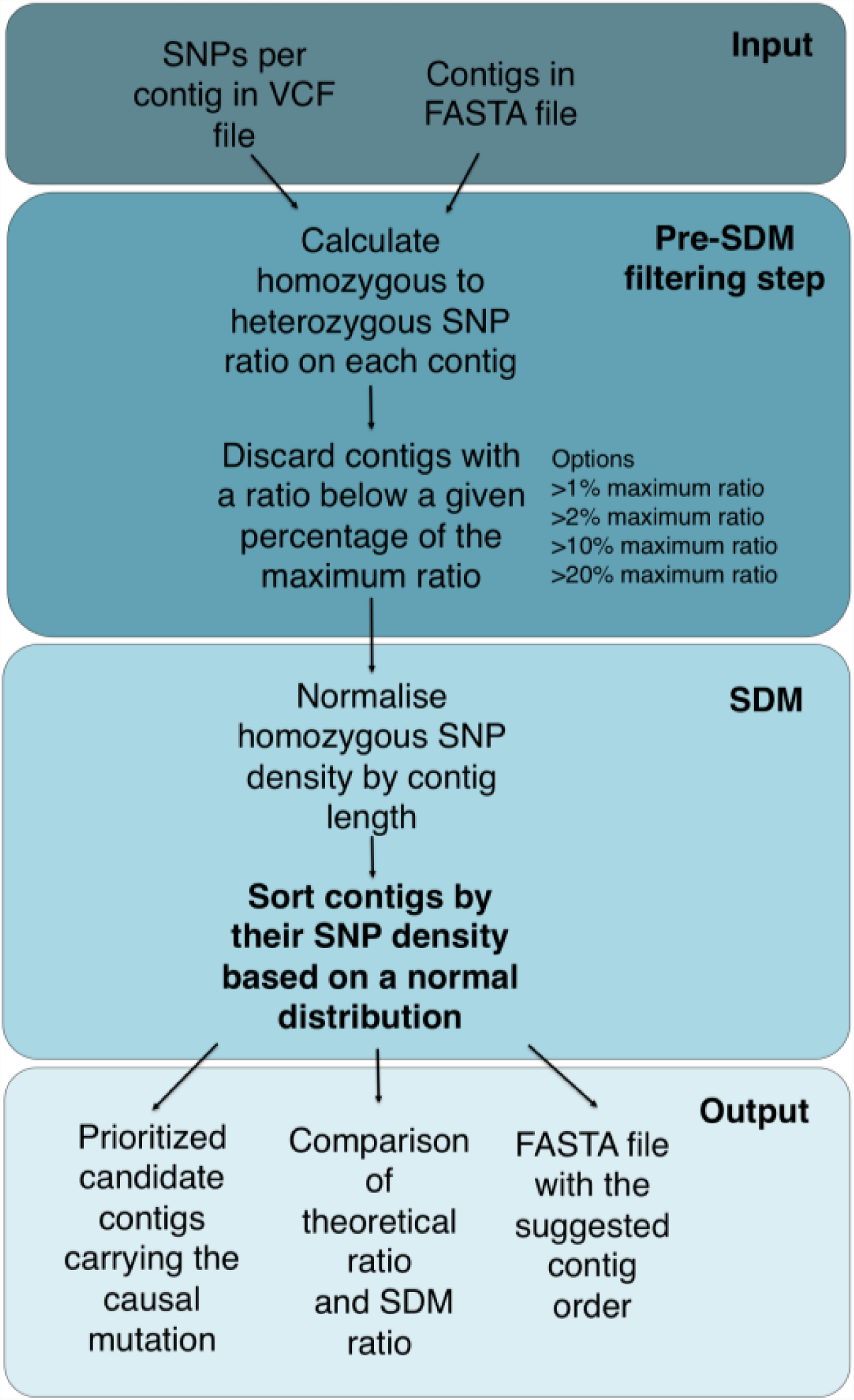
SDM workflow.

### 2.7. Parental filtering

To unmask the high homozygous SNP peak, we performed a filtering step to reduce the SNP density. The parental reads were also mapped to the *A. thaliana* reference genome as explained in section 2.6 followed by a step of SNP calling. The SNPs present in the non-mutant parental reads were not induced by the mutagen (EMS) and can be discarded from the mutant VCF file.

We ran manage_vcf.rb to filter the background SNPs. The workflow we followed can be found in the Supplementary File 2 and the protocol is available in the README file deposited at https://github.com/pilarcormo/SNP_distribution_method/tree/master/Reads.

### 2.8. Centromere removal

A great part of the variability observed in the genomes was due to the presence of centromeres. We ran remove_cent.rb to discard the SNP positions that were due to the centromere variability. The workflow used to filter the SNPs can be found in the Supplementary File 2 and a detailed protocol is available in the README file deposited at https://github.com/pilarcormo/SNP_distribution_method/tree/master/Reads.

### 2.9. SNP density analysis

We ran SNP_density.rb to take the absolute number of homozygous SNPs before and after filtering. The command instructions are available at the Supplementary File 2.

The output CSV file is available at https://github.com/pilarcormo/SNP_distribution_method/blob/master/Reads/density.csv. It shows the number of homozygous SNPs per chromosome and per forward genetic screen (BCF2, OCF2, sup#1, mob1, mob2). We obtained the total number of homozygous SNPs in the genome by adding together the values per chromosome. We then created new CSV files for the back-cross and the out-cross experiments. These are available at https://github.com/pilarcormo/SNP_distribution_method/blob/master/Reads/density_sum_back.csv and https://github.com/pilarcormo/SNP_distribution_method/blob/master/Reads/density_sum_out.csv.

We used the R code at https://github.com/pilarcormo/SNP_distribution_method/blob/master/R_scripts/SNP_filtering.R to plot the total number of homozygous SNPs before filtering, after parental filtering and after centromere removal. We also plot the number of candidate SNPs obtained after running SDM.

We plotted the homozygous and heterozygous SNP densities obtained after filtering for each study together with the ratio signal to identify the high density peaks in the distribution. The R code was deposited at https://github.com/pilarcormo/SNP_distribution_method/blob/master/Reads/filtering.md.

### 2.10. Probability plots

To analyse the correlation of the homozygous SNP density in forward genetic screens to a normal distribution, we created probability plots (QQ-plots). We used the homozygous SNP positions in the chromosome where the causative mutation was located. The R code is available at https://github.com/pilarcormo/SNP_distribution_method/blob/master/Reads/qqplot.md.

### 2.11. Analysis of average contig size in different whole genome assemblies

Plant genome assemblies at contig level from http://www.ncbi.nlm.nih.gov/assembly/organism/3193/all/ were used to define a more realistic contig size in our model genomes. We analysed contig assemblies from January 2013 to June 2015 which provided a full genome representation and a genome coverage higher than 1x. Only those providing the sequencing technology and the N50 contig size were selected to analyse the contig size distribution.

The table with the chosen assemblies and the results are available at https://github.com/pilarcormo/SNP_distribution_method/tree/master/Contigs.

We calculated the N50 density and the median of the distribution. We focused on the 16 assemblies built on Illumina Hiseq data and tried to define a model for the N50 contig size change over genome length. After applying logarithms, we first adjust a linear regression (lm) and then we apply a Generalised Additive Model (GAM) to fit non-parametric smoothers to the data without specifying a particular model. The R code can be found at https://github.com/pilarcormo/SNP_distribution_method/blob/master/Contigs/contigs.R.

### 2.12. Model genomes based on real SNP densities

We created new model genomes using the homozygous and heterozygous SNP densities obtained from the forward genetic screens after parental filtering and centromere removal. Three minimum contig sizes (2,000, 5,000 and 10,000 bp) were used, with maximum values of 4,000, 10,000 and 20,000 bp respectively. The SNP densities can be found at https://github.com/pilarcormo/SNP_distribution_method/tree/master/arabidopsis_datasets/SNP_densities.

The genomes were generated by running model_genome_real_hpc.rb. The command instructions are available in the Supplementary File 2.

The genomes generated are available at https://github.com/pilarcormo/SNP_distribution_method/tree/master/arabidopsis_datasets/No_centromere. They are classified by contig size.

### 2.13. SDM with real SNP densities

The model genomes generated as described in 2.12 were used to prove the efficiency of SDM to identify the genomic region carrying the causative mutation. The Ruby code for SDM is available at SNP_distribution_method_variation.rb. The input and output specification for SDM can be found in the README file in the main project Github repository https://github.com/pilarcormo/SNP_distribution_method.

Instead of specifying a percentage of the maximum ratio to filter the contigs, we used an automatic approach to tailor the threshold for each specific SNP density and contig length. The default percentage of the maximum ratio used was 1%. After the first filtering round, if the amount of discarded contigs is less than 3% of the original amount of contigs, the percentage of the maximum ratio is increased by 2 and the filtering is repeated until the specified condition is met.

The command instructions used to run SDM on the model genomes are available in Supplementary File 2.

## 3. Results and discussion

### 3.1. SDM is effective over a range of genome lengths and realistic fragment sizes

We created model genomes based on *Arabidopsis thaliana* chromosomes to develop our mutant identification method. Due to its relatively small and well-annotated genome, *Arabidopsis thaliana* is a widely used organism for forward genetic screening. SNP densities in several mapping-by-sequencing experiments in *Arabidopsis* are publicly available (see section 3.3 for examples) so they could be used as a starting point to develop our methodology.

By generating customised genomes we were able to rapidly alter different parameters such as genome length, contig size or SNP density to analyse their effect on the accuracy of the detection method. Our dynamic way of creating model genomes helped us define all the different aspects that should be taken into account when analysing the SNP distribution. The causative mutation was defined manually by us, so we could measure the shift between defined and predicted value. We generated the model genomes by assigning an idealised SNP distribution to a set of randomly shuffled sequences that imitate contigs assembled from HTS.

We first focused on small genomes with high SNP density as described in section 2.1. The SNP Distribution Method (SDM) sorts the sequence fragments by their SNP density values so that they follow an idealised normal distribution and then takes the highest density value as candidate. We measured the deviation from the expected peak (‘shift’ defined in section 2.2) and obtained consistent results for all the replicates (Fig. 2), so we can conclude that SDM works effectively over a range of different genome lengths and contig sizes.

**Figure 2.**
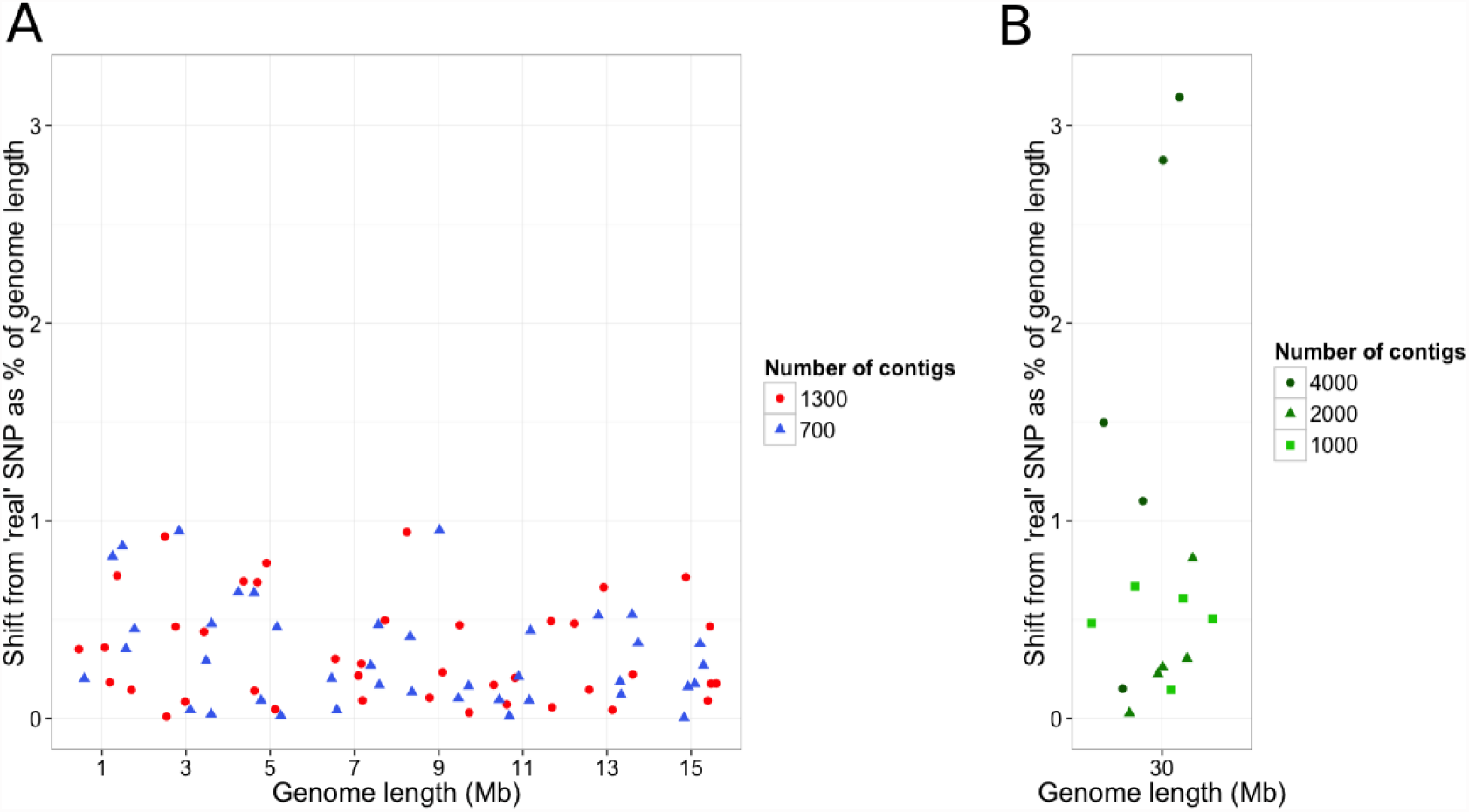
SDM shift from the expected causative mutation location in model genomes. 5 replicates of each model genome. **(A)** Model genome size ranges from 1 to 15 Mb with two contig sizes (1,300 contigs and 700). **(B)** Whole-sized *Arabidopsis thaliana* chromosome 1 model genomes with three contig sizes (4,000, 2,000 and 1,000 contigs)

The shift from the causative mutation assigned in the model was lower than 1% in the small SNP-rich genomes (Fig. 2A) and whole-sized genomes when they were fragmented in 1,000 and 2,000 contigs but not in those fragmented in 4,000 contigs (Fig. 2B), indicating that more contigs make the sorting step harder. Since we used a constant SNP density for all the contig sizes, SNPs are spread over shorter fragments so the high density peak is fragmented and the sorting becomes more complicated. We observed a slight decrease in SDM efficiency when the average contig size for a 30 Mb genome is 7,500 bp, i.e 4,000 contigs (Fig. 2B).

### 3.2 A pre-filtering step based on the homozygous to heterozygous SNPs ratio improves SDM accuracy

SDM was able to identify the high density peak in model genomes when the idealised causative mutation was located in the middle of the distribution, as the number of fragments at both tails of the distribution was the same. High sensitivity is only achieved in the high SNP density area (peak of the distribution) while the contigs located in the tails cannot be sorted by their SNP density. When we shifted the causal mutation in our model to one side (one tail was longer than the other), SDM was not able to sort the contigs in the tails properly. Even though the contigs located in the peak were correct, the algorithm was not able to sort the low SNP density regions and the highest kernel density value in the distribution shifted from the previously defined point (Fig. 3A).

**Figure 3.**
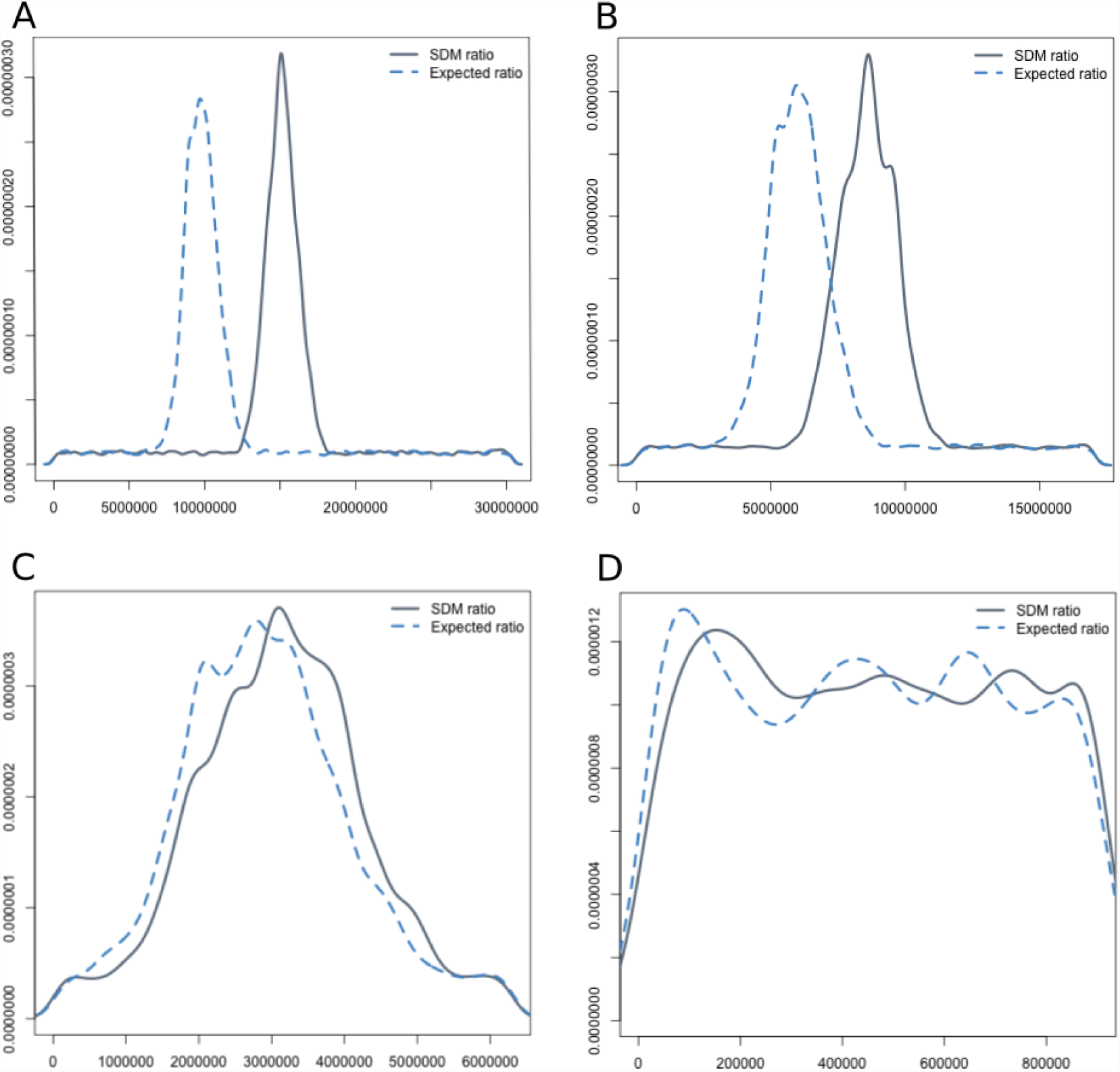
Effect of a pre-filtering step based on the homozygous to heterozygous SNP ratio in model genomes. The ratio is calculated per contig and those contigs falling below a given percentage of the maximum ratio are discarded. The expected ratio was measured in the correctly ordered fragments. The SDM ratio was measured after SDM sorting. **(A)** Threshold = 1% of the maximum ratio. **(B)** Threshold = 5% of the maximum ratio. **(C)** Threshold = 10% of the maximum ratio. **(D)** Threshold = 20% of the maximum ratio.

We used a threshold value based on the ratio of homozygous to heterozygous SNPs to discard contigs located furthest away from the causative mutation. We excluded those contigs with a ratio below a given percentage of the maximum ratio. Only those contigs in the region of interest are sorted and we can assess the contigs in which the mutation is to be found, dismissing an uninformative part of the genome. To test the influence of the threshold we shifted the mean in the normal distribution 20% as described in section 2.1. With a high SNP density (1 SNP every 3000 bp) and a standard deviation of 1 Mb, the ratio peak matched the expected peak when 10% of the maximum ratio was used as a threshold (Fig. 3C), and approximately a 20% of the genome was discarded.

### 3.3 Filtering background SNPs and centromeres unmasks the high homozygous SNP density peak in bulked segregant analysis in Arabidopsis

We selected different datasets of bulked segregant analysis of a mutation segregating in an out-crossed [17, 18] or back-crossed [19, 20] population and performed conventional genome alignment and variant calling.

In the out-cross experiment OCF2, Galvão et al identified a mutation causing late flowering on the *SOC1* gene (2: 18807538..18811047) [17]. Allen at al analysed the mutant individuals showing leaf hyponasty to identify a gene involved in the *Arabidopsis* microRNA pathway (BCF2) [19] and identified the causal SNP in *HASTY* (3: 1401271..1408197). In the forward screen done in the immune-deficient *bak1-5* background to identify new components involved in plant immunity, Monaghan et al found 2 causative mutations in the gene encoding the calcium-dependent protein kinase CPK28 (5: 26456285..26459631) for both *bak1-5 mob1* and *bak1-5 mob2* [20]. Uchida et al identified the sup#1 mutation on the *SGT1b* gene (4: 6851277..6853860) [18]. The techniques used to identify the mutations were different in every case (Table 1).

We analysed the total number of homozygous SNPs (Fig. 4). When the mutant individual is out-crossed to a distant mapping line (OCF2 and sup#1), the SNP density is up to 20 times higher than in the case of back-crossing to the parental line (BCF2 and mob mutants). In the back-crossed populations we identified around 1,700 homozygous SNPs in the whole genome resulting in an overall density of ∼1 SNP every 70 kb (Table 2). We identified 9,208 homozygous SNPs in the first out-crossed population (OCF2) and 27,578 SNPs in the second out-crossed population (sup#1). The overall density was 1 SNP in every 12.9 kb for OCF2 and 1 SNP in every 4.3 kb for sup#1 (Table 2).

**Table 2.** Total number of homozygous SNPs identified in each forward genetic screen, measurement of the homozygous SNP density correlation to a theoretical normal distribution after parental filtering and centromere removal and standard deviation (sd) of the normal distribution.

| Sample | Total hm SNPs | $r^2$ | sd (Mb) |
| --- | --- | --- | --- |
| OCF2 | 9208 | 0.94 | 6.01 |
| BCF2 | 1678 | 0.95 | 3.20 |
| mob1 | 1694 | 0.94 | 7.20 |
| mob2 | 1726 | 0.92 | 5.79 |
| sup#1 | 27548 | 0.98 | 3.72 |

**Figure 4.**
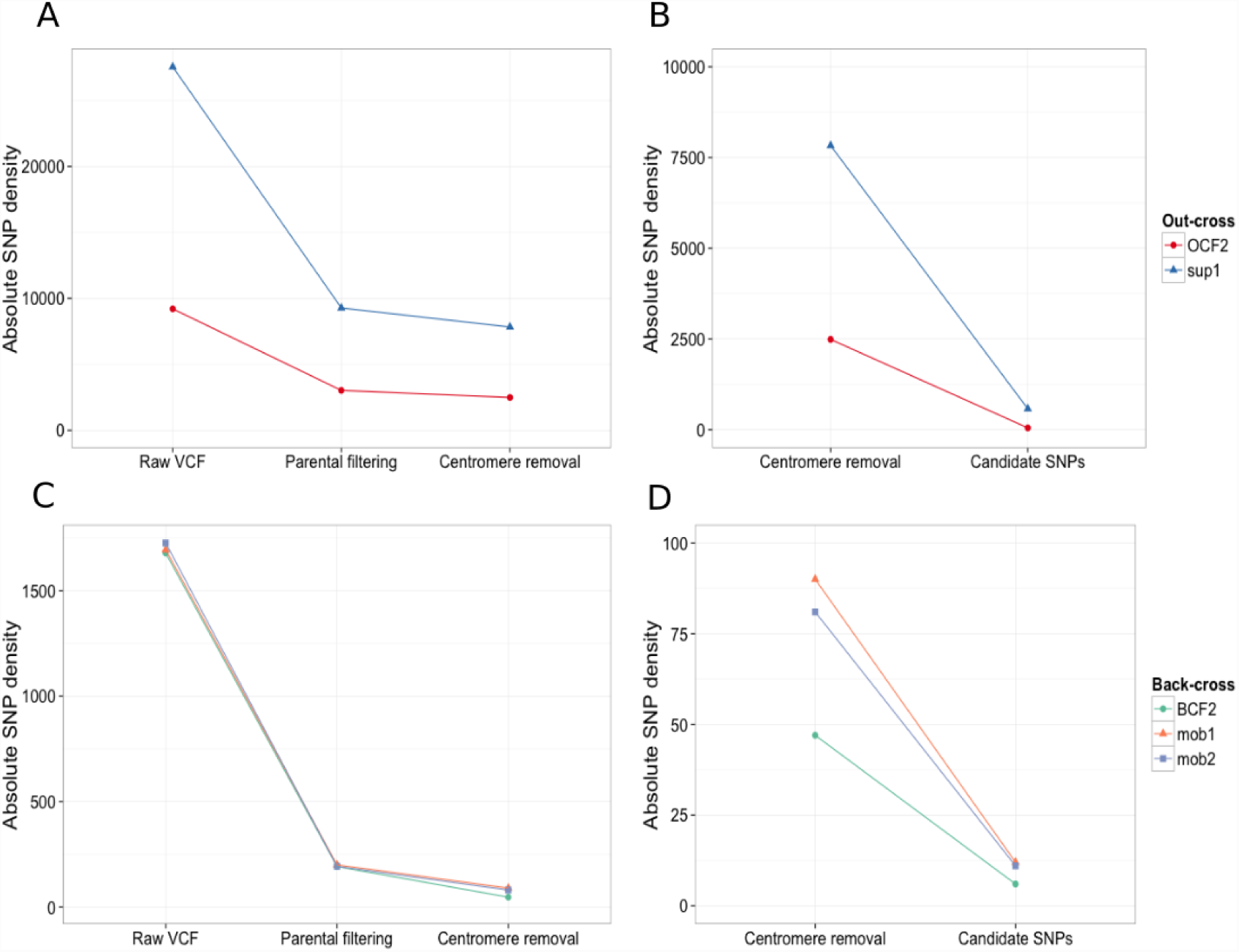
Absolute number of homozygous SNPs before and after filtering in independent **(A)** back-crossed and **(B)** out-crossed populations. The final candidate positions after running SDM were also compared in the **(C)** back-crossed and **(D)** out-crossed populations.

Parental filtering was very effective to reduce the SNP density and unmask the SNP linkage around the causative mutation especially in the out-crossed populations where the starting density was higher. The SNPs present in the non-mutant parental reads were filtered from the mutant SNP lists. In the back-cross studies, the absolute homozygous SNP number was reduced up to 1/9 of the original amount (Fig. 4A) after parental filtering. The total number of homozygous SNPs was reduced to 1/3 of the original amount in out-crossed populations (Fig. 4C). Even though the centromere removal did not reduce the total number of SNPs in the same proportion as parental filtering did, it was essential to unhide the normal distribution around the causative mutation. The centromere is characterised by high repeat abundance (often >10,000 copies per chromosome) [27], so high variability in a few hundred bp region generates a high SNP density peak which obscures the peak of interest.

The main advantage of working with already identified mutations is the ability to focus on the chromosome on which the mutation was previously described. We identified a unique peak in the area where the causative mutation was described when we plotted the homozygous SNP density obtained after filtering. We calculated the homozygous to heterozygous ratio for each contig and the ratio values were overlapped to the SNP densities. Fig. 5A shows the density plots obtained for the back-crossed populations and Fig. 5B shows the density plots for the out-crossed populations. The location of the causative mutation previously described is always within the high SNP density peak [17, 18, 19, 20].

**Figure 5.**
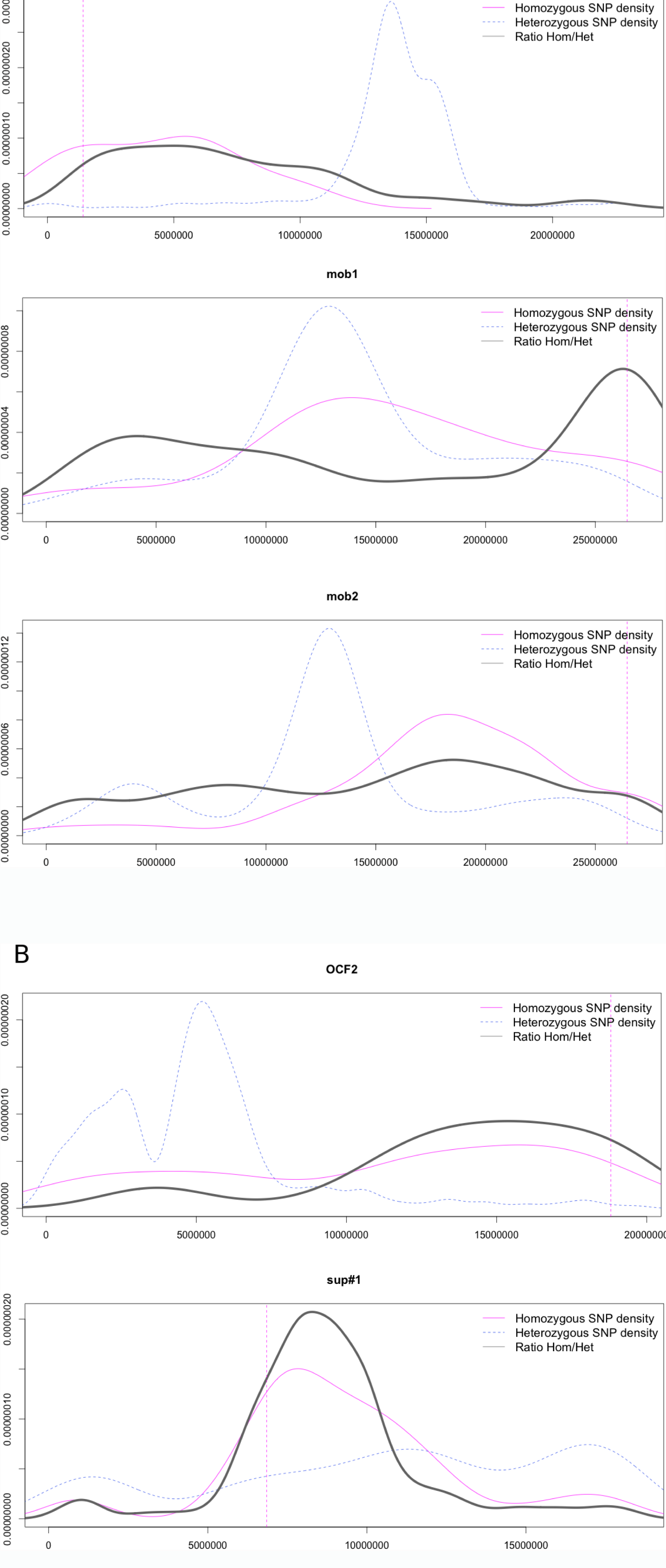
Identification of high homozygous SNP density peaks surrounding the causal mutation in 5 independent studies. Overlapping homozygous and heterozygous SNP densities and hom/het ratios for OCF2, BCF2, bak1-5 mob1/mob2 and sup#1.

### 3.4. Homozygous SNPs in forward genetic screens are normally distributed around the causative mutation

We observed a unique peak in the SNP distribution around the causative mutation for all the samples (Fig. 5). The next step was to analyse the correlation of the SNP density to a theoretical probability distribution. We created probability plots (sometimes called QQ-plots) with the homozygous SNP densities in the back-crossed and out-crossed populations (Fig. 6) before filtering, after parental filtering and after centromere removal. The filtering steps reduced the complexity of the distribution (**Additional Figure 2**) and improved the degree of correlation to a normal distribution (Fig. 6) as proven in the r^2^ increase.

**Figure 6.**
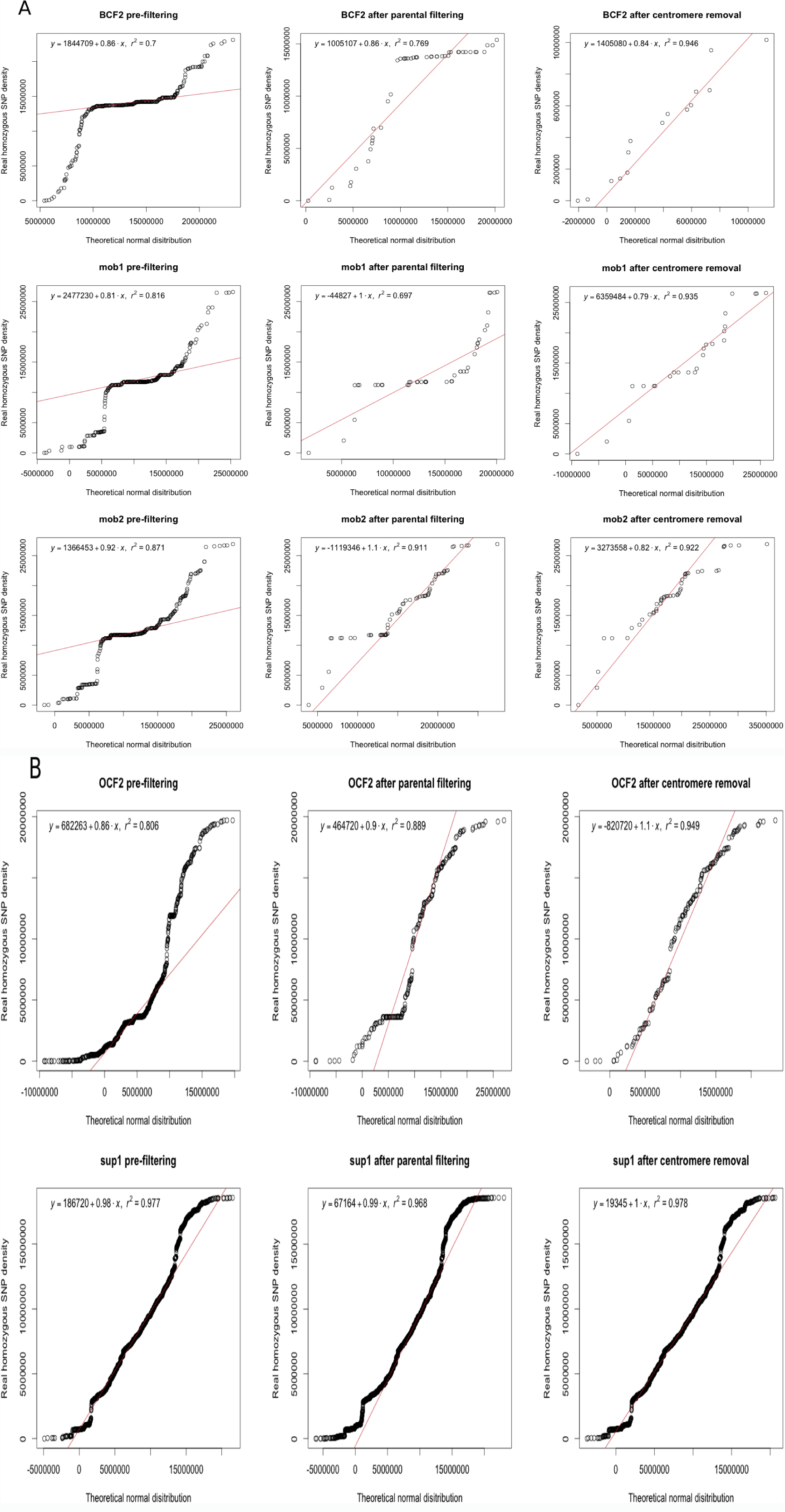
Measurement of the homozygous SNP density correlation to a normal distribution in back-crossed and out-crossed populations by probability (QQ) plots before filtering, after parental filtering and after centromere removal. Simple linear regression was used to verify the correlation.

The idea of removing non-unique SNPs is not new, and all the mapping-by-sequencing studies we analysed did the same filtering to some extent.

Our results indicate a good correlation between the homozygous SNP frequencies after filtering and a normal distribution. We further validated the correlation by a simple linear regression (r^2^ ≥ 0.92). The standard deviation was between 3 and 7 Mb (Table 2).

### 3.5. The N50 contig size in plant genome assemblies depends on genome size and sequencing technology

To obtain parameters for the generation of realistic model genomes, we analysed 29 assemblies at contig level. The relationship between genome length and N50 contig size was not strong and other aspects such us the sequencing technology used (Fig. 7A) or the genome coverage had a high impact on the final N50 contig size. The median value of the N50 contig length for all the 29 assemblies is 11,517 bp while it is reduced to 5,484 bp when analysing only assemblies from Illumina HiSeq data (Fig. 7B). To decrease the effect of the technology used, we focused on the 16 assemblies built with Illumina Hiseq as it was the most popular sequencing strategy. Then we tried to establish a model that could explain the N50 change over genome length.

**Figure 7.**
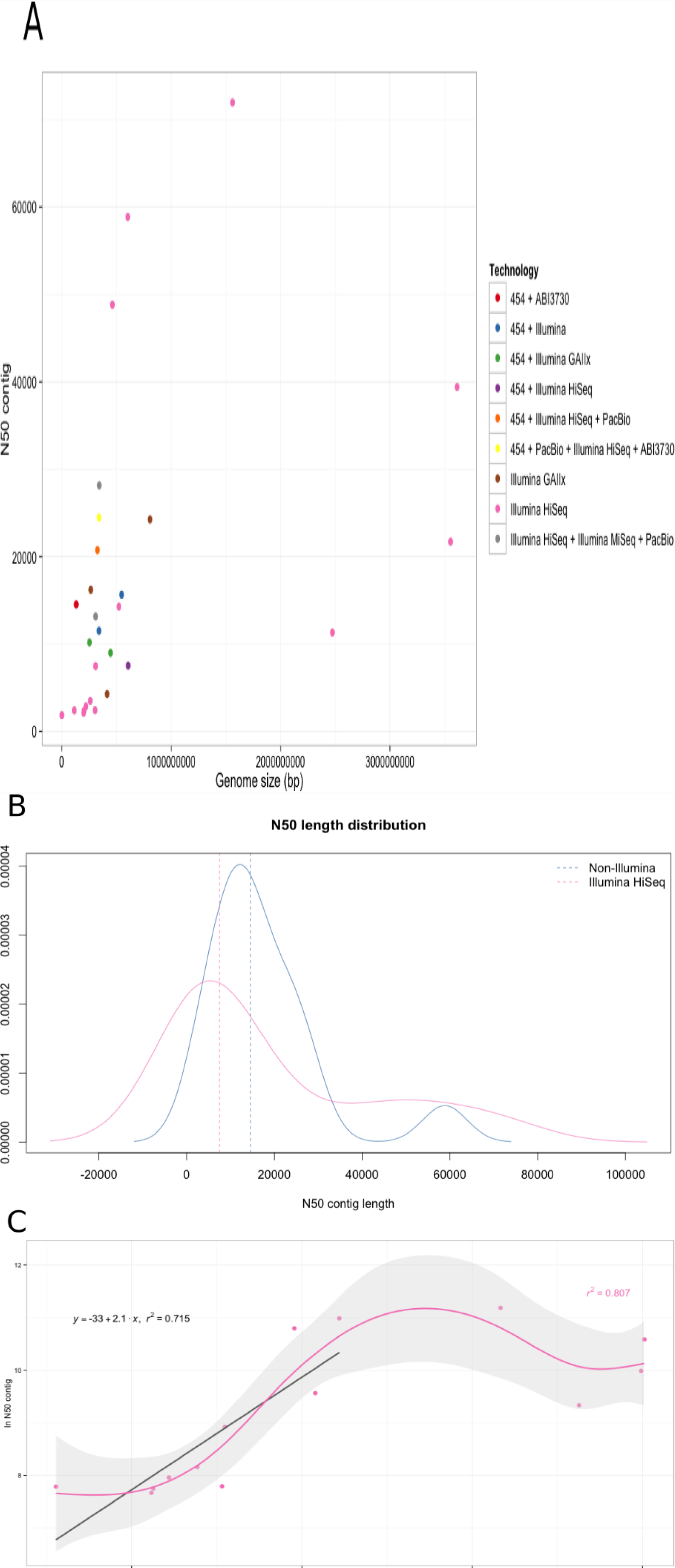
Analysis of average contig size in different genome assemblies. **(A)** N50 contig size vs Genome size in 29 whole genome assemblies at contig level. Sequencing technology or the combination of sequencing technologies is colour coded. **(B)** N50 contig size distribution for Illumina HiSeq assemblies (pink) and for assemblies from other sequencing technologies (blue). Medians are represented by the dashed lines. **(C)** Model for the non-linear relationship between N50 contig size and genome size in Illumina HiSeq assemblies.

There was not a direct mechanism to fit a model to the data. We could identify a linear correlation (r^2^ = 0.72) between N50 size and genome length when genome size was below 1 Gb (Fig. 7C). For larger genomes the linear relationship was not maintained. We also applied a Generalised Additive Model (GAM) to fit non-parametric smoothers to the data (r^2^ = 0.81) (Fig. 7C).

We used 3 different contig sizes to create the model genomes. The first two model genomes were built using the N50 median values. We chose 10,000 bp (based on the median for all the assemblies) and 5,000 bp (based on the median for Illumina Hiseq data) (Fig. 7B). The smallest contig size was decided looking at the linear model defined for Illumina HiSeq assemblies (Fig. 7C). The minimum contig size decided for these model genomes was 2,000 bp.

### 3.6. SDM identifies the genomic region carrying the causal mutation previously described by other methods

We used the SNP densities obtained from OCF2, BCF2, mob1, mob2 and sup#1 datasets after parental filtering and centromere removal to build new model genomes. We chose the chromosomes from *Arabidopsis thaliana* in which the mutations were described (Table 1) and split them into fragments of size specified in section 3.5. The SNP density used to build the model genomes was the same for all the contig sizes.

We regained the normal distribution for all the datasets after shuffling the contig order and running SDM. The results for all the model genomes generated were deposited in a Github repository at https://github.com/pilarcormo/SNP_distribution_method/tree/master/arabidopsis_datasets/No_centromere.

Our prior knowledge about the correct contig order allowed us to define the real chromosomal positions in the artificial contigs identified by SDM as candidates. In that way, we were able to adjust the method to maximise its efficiency (Table 3). Contig size had an effect on the number of candidate contigs provided by SDM. When the minimum contig sizes were 2 and 5 kb, the SNP positions were split into different contigs and it was harder for SDM to find the correct contig order. As a result, when the average contig size was below 10 kb, 20 candidate contigs were needed for back-crossed populations and 40 for out-crossed populations due to the high SNP densities. When the average contig size was greater than 10 kb, 12 candidate contigs in the middle of the distribution were enough to contain the causative mutation. The candidate region ranges from 60 to 180 kb depending on the contig size and the type of cross.

**Table 3.** Results of SDM mutant identification when an automatic filtering approach is used to discard contigs. Three different contig sizes were analysed and the percentages of the maximum ratio used as threshold are specified. The number of discarded contigs is in brackets (discarded contigs/total).

| Sample | Contig size (kb) | Threshold | Identification |
| --- | --- | --- | --- |
| OCF2 | 2-4 | 5% (230/6568) | **Unsuccessful** |
|  | 5-10 | 3% (189/2634) | Successful |
|  | 10-20 | 3% (186/1328) | Successful |
| BCF2 | 2-4 | 35% (7807/7821) | Successful |
|  | 5-10 | 21% (108/3130) | Successful |
|  | 10-20 | 21% (95/1562) | Successful |
| mob1 | 2-4 | 17% (254/8992) | **Unsuccessful** |
|  | 5-10 | 15% (239/3603) | Successful |
|  | 10-20 | 15% (220/1805) | Successful |
| mob2 | 2-4 | 35% (8950/8994) | Successful |
|  | 5-10 | 21% (195/3582) | Successful |
|  | 10-20 | 21% (189/1804) | Successful |
| sup#1 | 2-4 | 3% (228/6201) | Successful |
|  | 5-10 | 3% (153/2491) | Successful |
|  | 10-20 | 3% (93/1240) | Successful |

We could not define a universal cut-off value based on the homozygous to heterozygous ratio for all the different SNP densities and crosses, as sometimes the region with a high ratio was narrow due to a high SNP linkage in the area, while in other cases, the increase in the ratio was progressive, and the peak was wider. When we worked with real densities, we used an automatic approach that tailors the threshold for each specific SNP density and contig length as explained in section 2.13. Table 3 shows the tailored thresholds and total discarded contigs for all the datasets.

We conclude that SDM is a rapid method to perform bulked segregant linkage analysis from back-crossed and out-crossed populations without relying on the availability of a reference genome. It is especially effective on contig sizes over 5 kb. Even though it can be accurate with smaller contigs, we defined the detection limit on 2-4 kb contigs. In 2 out of the 5 models, the contig carrying the candidate mutation was lost when the contig size oscillated between 2 and 4 kb.

## 4. Conclusions

Forward genetic screens are very useful to identify genes responsible for particular phenotypes. Due to the advances in HTS technologies, mutant genome sequencing has become quick and inexpensive. Mapping-by-sequencing methods available present certain limitations, complicating the mutation identification especially in non-sequenced species. To target this problem we proposed a fast, reference genome independent method to identify causative mutations. We showed that homozygous SNPs are roughly normally distributed in the mutant genome of back-crossed and out-crossed individuals. Based on that idea we defined a theoretical SNP distribution used by SDM to identify the genomic region where the causative mutation was located.

By using customised model genomes we could rapidly alter different parameters to tune the detection method. SDM was optimised for *Arabidopsis thaliana* and it was able to identify the contigs carrying the causative SNPs in four independent forward genetic screens. We conclude that SDM is especially successful at identifing the genomic region carrying a mutation in typical SNP densities when the contig size is at least 2 kb. A raise in the SNP density in out-crossed experiments increased the number of candidate contigs, but SDM was able to predict the contig containing the causal mutation. SDM is a promising method for causative mutation identification and we hope to reproduce the good results obtained for *Arabidopsis thaliana* in other organisms. We aim to apply SDM in forward genetic screens of species where a reference genome is not yet available.

## Acknowledgements

We thank Martin Page, Ghanasyam Rallapalli and Christian Schudoma for technical assistance and valuable discussions. We thank Christian Schudoma for the code refactoring sessions and for carefully reading the manuscript.

## References

1. Page DR, Grossniklaus U: The art and design of genetic screens:*Arabidopsis thaliana*. Nat Rev Genet 2002, 3:124–36.

2. Etherington GJ, Monaghan J, Zipfel C, MacLean D: Mapping mutations in plant genomes with the user-friendly web application CandiSNP. Plant Methods 2014, 10:41.

3. Schneeberger K: Using next-generation sequencing to isolate mutant genes from forward genetic screens. Nat Rev Genet 2014, 15:662–76.

4. Michelmore RW, Paran I, Kesseli RV: Identification of markers linked to disease-resistance genes by bulked segregant analysis: A rapid method to detect markers in specific genomic regions by using segregating populations. Proc Natl Acad Sci U S A 1991, 88:9828–32.

5. Schneeberger K, Ossowski S, Lanz C, Juul T, Petersen AH, Nielsen KL, Jørgensen J-E, Weigel D, Andersen SU: SHOREmap: Simultaneous mapping and mutation identification by deep sequencing. Nat Methods 2009, 6:550–1.

6. Sun H, Schneeberger K: SHOREmap v3.0: Fast and accurate identiffcation of causal mutations from forward genetic screens. Methods Mol Biol 2015, 1284:381–95.

7. Austin RS, Vidaurre D, Stamatiou G, Breit R, Provart NJ, Bonetta D, Zhang J, Fung P, Gong Y, Wang PW, McCourt P, Guttman DS: Next-generation mapping of arabidopsis genes. Plant J 2011, 67:715–25.

8. Wurtzel O, Dori-Bachash M, Pietrokovski S, Jurkevitch E, Sorek R: Mutation detection with next-generation resequencing through a mediator genome. PLoS One 2010, 5:e15628.

9. Livaja M, Wang Y, Wieckhorst S, Haseneyer G, Seidel M, Hahn V, Knapp SJ, Taudien S, Schön C-C, Bauer E: BSTA: A targeted approach combines bulked segregant analysis with next-generation sequencing and de novo transcriptome assembly for sNP discovery in sunflower. BMC Genomics 2013, 14:628.

10. Nordström KJV, Albani MC, James GV, Gutjahr C, Hartwig B, Turck F, Paszkowski U, Coupland G, Schneeberger K: Mutation identiffcation by direct comparison of whole-genome sequencing data from mutant and wild-type individuals using k-mers. Nat Biotechnol 2013, 31:325–30.

11. Song J, Bradeen JM, Naess SK, Raasch JA, Wielgus SM, Haberlach GT, Liu J, Kuang H, Austin-Phillips S, Buell CR, Helgeson JP, Jiang J: Gene RB cloned from *Solanum bulbocastanum* confers broad spectrum resistance to potato late blight. Proc Natl Acad Sci U S A 2003, 100:9128–33.

12. Iqbal Z, Caccamo M, Turner I, Flicek P, McVean G: De novo assembly and genotyping of variants using colored de bruijn graphs. Nat Genet 2012, 44:226–32.

13. Minevich G, Park DS, Blankenberg D, Poole RJ, Hobert O: CloudMap: A cloud-based pipeline for analysis of mutant genome sequences. Genetics 2012, 192:1249–69.

14. Abe A, Kosugi S, Yoshida K, Natsume S, Takagi H, Kanzaki H, Matsumura H, Yoshida K, Mitsuoka C, Tamiru M, Innan H, Cano L, Kamoun S, Terauchi R: Genome sequencing reveals agronomically important loci in rice using MutMap. Nat Biotechnol 2012, 30:174–8.

15. Takagi H, Tamiru M, Abe A, Yoshida K, Uemura A, Yaegashi H, Obara T, Oikawa K, Utsushi H, Kanzaki E, Mitsuoka C, Natsume S, Kosugi S, Kanzaki H, Matsumura H, Urasaki N, Kamoun S, Terauchi R: MutMap accelerates breeding of a salt-tolerant rice cultivar. Nat Biotechnol 2015, 33:445–9.

16. Lamesch P, Berardini TZ, Li D, Swarbreck D, Wilks C, Sasidharan R, Muller R, Dreher K, Alexander DL, Garcia-Hernandez M, Karthikeyan AS, Lee CH, Nelson WD, Ploetz L, Singh S, Wensel A, Huala E: The arabidopsis information resource (TAIR): Improved gene annotation and new tools. Nucleic Acids Res 2012, 40(Database issue):D1202–10.

17. Galvão VC, Nordström KJV, Lanz C, Sulz P, Mathieu J, Posé D, Schmid M, Weigel D, Schneeberger K: Synteny-based mapping-by-sequencing enabled by targeted enrichment. Plant J 2012, 71:517–26.

18. Uchida N, Sakamoto T, Tasaka M, Kurata T: Identification of EMS-induced causal mutations in arabidopsis thaliana by next-generation sequencing. Methods Mol Biol 2014, 1062:259–70.

19. Allen RS, Nakasugi K, Doran RL, Millar AA, Waterhouse PM: Facile mutant identiffcation via a single parental backcross method and application of whole genome sequencing based mapping pipelines. Front Plant Sci 2013, 4:362.

20. Monaghan J, Matschi S, Shorinola O, Rovenich H, Matei A, Segonzac C, Malinovsky FG, Rathjen JP, MacLean D, Romeis T, Zipfel C: The calcium-dependent protein kinase CPK28 buffers plant immunity and regulates BIK1 turnover. Cell Host Microbe 2014, 16:605–15.

21. DePristo MA, Banks E, Poplin R, Garimella KV, Maguire JR, Hartl C, Philippakis AA, Angel G del, Rivas MA, Hanna M, McKenna A, Fennell TJ, Kernytsky AM, Sivachenko AY, Cibulskis K, Gabriel SB, Altshuler D, Daly MJ: A framework for variation discovery and genotyping using next-generation DNA sequencing data. Nat Genet 2011, 43:491–8.

22. Li H, Handsaker B, Wysoker A, Fennell T, Ruan J, Homer N, Marth G, Abecasis G, Durbin R, 1000 Genome Project Data Processing Subgroup: The sequence alignment/Map format and sAMtools. Bioinformatics 2009, 25:2078–9.

23. Bolger AM, Lohse M, Usadel B: Trimmomatic: A flexible trimmer for illumina sequence data. Bioinformatics 2014, 30:2114–20.

24. Li H, Durbin R: Fast and accurate long-read alignment with burrowswheeler transform. Bioinformatics 2010, 26:589–95.

25. Koboldt DC, Zhang Q, Larson DE, Shen D, McLellan MD, Lin L, Miller CA, Mardis ER, Ding L,Wilson RK: VarScan 2: Somatic mutation and copy number alteration discovery in Cancer by exome sequencing. Genome Res 2012, 22:568–76.

26. Koboldt DC, Chen K, Wylie T, Larson DE, McLellan MD, Mardis ER, Weinstock GM, Wilson RK, Ding L: VarScan: Variant detection in massively parallel sequencing of individual and pooled samples. Bioinformatics 2009, 25:2283–5.

27. Melters DP, Bradnam KR, Young HA, Telis N, May MR, Ruby JG, Sebra R, Peluso P, Eid J, Rank D, Garcia JF, DeRisi JL, Smith T, Tobias C, Ross-Ibarra J, Korf I, Chan SWL: Comparative analysis of tandem repeats from hundreds of species reveals unique insights into centromere evolution. Genome Biol 2013, 14:R10.

